# Hydrogel microneedle patch for low-cost, colorimetric, continuous glucose monitoring

**DOI:** 10.64898/2026.10.12.758503

**Authors:** Anthony Du, Hannah Newman, Julian Diaz-Ayala, Mason Doyle, Sneha Sengupta, Yongjae Lee, Emma Chory, Cameron M. Kim

**Affiliations:** Department of Biomedical Engineering, Duke University, Durham, NC 27705, USA

## Abstract

Diabetes is a chronic condition characterized by dysregulated blood glucose levels. Continuous glucose monitors (CGMs) enable minimally invasive, real-time monitoring of glucose levels, but broader access requires devices that are both effective and affordable. Popular CGMs such as Dexcom and FreeStyle Libre have high accuracy and are integrated with mobile applications to display real-time data and alerts but are limited by high cost. We developed a low-cost, electronic-free, microneedle hydrogel patch CGM that utilizes colorimetric detection to continuously monitor glucose levels. The proposed patch penetrates the skin and absorbs interstitial fluid (ISF) containing glucose. This glucose reacts with glucose oxidase (GOx) enzyme embedded in the hydrogel, producing gluconic acid and decreasing the pH of the hydrogel. As glucose levels increase, decreasing pH levels are reported by the loaded pH indicator bromocresol purple (BCP) via a color change from purple to yellow. This system was validated through a combination of *in vitro* experiments and COMSOL computational modeling to assess performance at physiologically relevant glucose concentrations. The resulting hydrogels demonstrated reversible and dynamic color changes in response to glucose levels ranging from 3.5-10 mmol/L (63-180 mg/dL), supporting their potential for continuous monitoring applications. A quantitative, image-based analysis method enabled estimation of glucose concentration from photographs, demonstrating strong predictive performance (RMSE=8.59 mg/dL, R^2^=0.94). Although optimization of mechanical strength and diffusion kinetics remain necessary, these findings highlight the promise of integrating colorimetric sensing with microneedle-based sampling for affordable, user-friendly glucose monitoring.

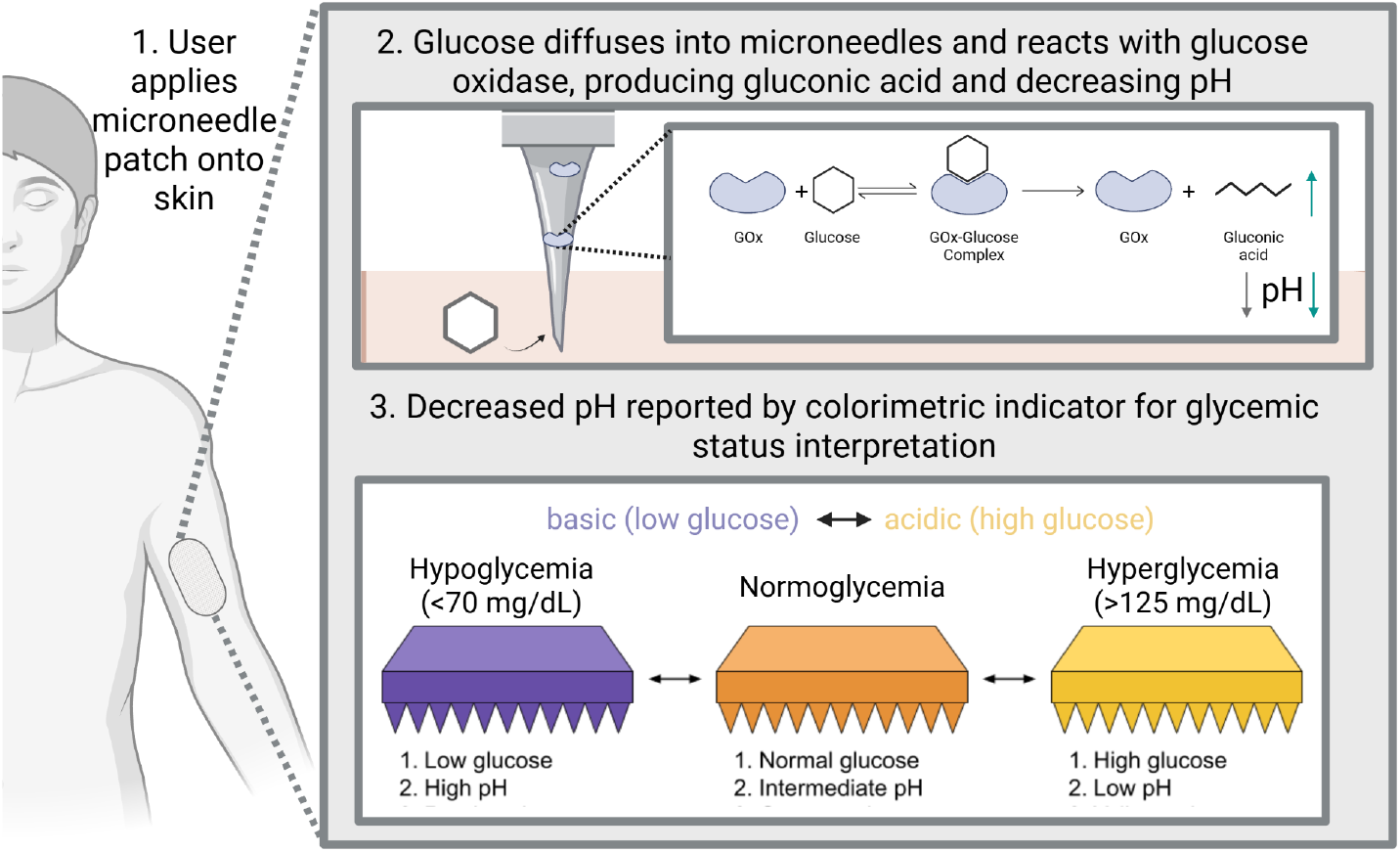

## INTRODUCTION

Diabetes is a chronic condition caused by the impairment or lack of insulin signaling which results in poor glycemic control and affects 830 million people worldwide (NCD Risk Factor Collaboration (NCD-RisC), 2024). Diabetes can lead to hyperglycemia or hypoglycemia, defined as high (>125 mg/dL) or low (<70 mg/dL) blood sugar, respectively (Antar et al., 2023; Mathew & Thoppil, 2025; Mouri & Badireddy, 2025). Left untreated, diabetes can pose serious health issues, with associated complications including disease of the hepatic, renal, cardiovascular, and nervous systems, among others (Antar et al., 2023). Managing blood glucose levels typically requires daily injections of basal, slow-acting insulin along with bolus, fast-acting insulin before meals to avoid blood sugar spikes. Consistent blood glucose monitoring allows for timely adjustment of insulin dosing, resulting in greater time in healthy glucose level ranges and significantly decreased risk of complications (Holman et al., 2008; Lind et al., 2024).

There are a variety of methods for measuring glucose levels, each with unique limitations. The most common method relies on the oxidation of glucose by glucose oxidase (GOx), which produces gluconic acid and hydrogen peroxide (H_2_O_2_) in the presence of oxygen as the electron acceptor. Typically, H_2_O_2_ is reacted with a metal sensor to release electrons, which can be quantitatively detected via an electrical current and used to calculate glucose concentration (Witkowska Nery et al., 2016). However, such devices require power and expensive electrical components to function.

The gold standard clinical device for glucose testing, the YSI 2900C Biochemistry Analyzer, implements GOx chemistry through an immobilized enzyme layer housed in a benchtop device. The YSI 2900C can measure glucose levels to an accuracy of 2% (2.0 mg/dL), making it highly effective as a diagnostic (Food and Drug Administration. YSI 2900C Biochemistry Analyzer: 510(k) Substantial Equivalence Determination Decision Summary, n.d.). However, this device can cost around $12,000, limiting its use to the clinical setting (Han et al., 2020).

Finger-stick tests were developed to provide a cost effective and convenient alternative. Finger sticks monitor blood glucose with a drop of blood that the user collects from their finger using a small needle. While these glucometers typically cost ∼$15, accuracy can be as low as 32% (Olansky & Kennedy, 2010). These tests also only measure blood glucose at a single time point and may need to be administered up to 10 times a day (American Diabetes Association Professional Practice Committee, 2025). Moreover, finger-stick tests are invasive and painful, decreasing patient adherence and resulting in increased risk of complications (Egede et al., 2014; Lin et al., 2023).

Continuous glucose monitor (CGM) systems have revolutionized diabetes care by providing continuous, minimally invasive readings that allow patients to manage their condition more effectively. CGMs measure glucose levels in interstitial fluid (ISF) either through direct sampling or reverse iontophoresis every one to five minutes (Almurashi et al., 2023). Glucose levels in ISF correlate to plasma glucose levels with an average lag time of 8 to 10 minutes, which is accounted for in CGM software programs (Cengiz & Tamborlane, 2009). CGMs use real-time transmitters to update data to either a receiver or a smartphone, providing continuous insight into glycemic trends and automatically alerting patients of hypo- or hyperglycemic episodes (Olczuk & Priefer, 2018). Nonetheless, out of 2,600 American patients with diabetes, one in every five reported not using a CGM due to financial difficulties (Engler et al., 2018). Lack of access to CGMs may result in decreased glycemic control and increased rate of hospitalization due to hypo- and hyperglycemic events (Reaven et al., 2023).

Recent advances in hydrogel-based biosensing offer a pathway toward inexpensive CGM solutions that address these limitations. Hydrogels are polymeric networks that swell to absorb large amounts of water. Their high water content, soft structure, and porosity closely mimic human tissue (Ho et al., 2022), making hydrogels highly biocompatible, minimizing immune responses, and enabling easy physiological integration. Specifically, hydrogels can be used to form microneedles (MNs) which penetrate the skin and swell to absorb ISF (Aroche et al., 2025). Poly(ethylene glycol) diacrylate (PEGDA) hydrogels offer advantages due to their ability to retain large amounts of liquid upon swelling, making them ideal for absorbing ISF (Buxton et al., 2007). This also facilitates bidirectional diffusion of the molecular contents to allow for enzymatic reactions (Hakim Khalili et al., 2023).

Recent research has explored the potential for MN hydrogel CGMs. Previous work by Zeng et al. incorporated GOx, horseradish peroxidase (HRP), and 3,3′,5,5′-tetramethylbenzidine (TMB) into a MN hydrogel. The incorporation of GOx enabled the production of H_2_O_2_ from glucose diffusing into the patch. This was detected through HRP-catalyzed oxidation of TMB, producing a visible color change proportional to glucose concentration (Zeng et al., 2023). Other research from Wang et al. incorporated GOx into a responsive MN hydrogel. In this design, pH changes resulting from the production of gluconic acid modulated hydrogel swelling, allowing for quantitative readouts of glucose levels based on MN swelling ratios (Wang et al., 2023).

While these studies underscore the strong potential of hydrogel-based materials for cost-effective CGMs, they have a few key weaknesses preventing broader adoption of hydrogel-based CGMs. Zeng et al. used an approach that required a secondary enzyme, increasing the complexity and cost of the CGM. Moreover, the swelling-based readout used by Wang et al. requires physical measurement rather than simple visual inspection.

There remains a need for a simple, low-cost hydrogel patch CGM that utilizes colorimetric glucose detection. Color-based diagnostics have been adopted in other fields due to their ease of use and minimal instrumentation requirements. For example, lateral flow assays, such as home pregnancy and rapid infectious disease tests, have enabled point-of-care diagnostics with improved accessibility and user compliance (Kinyua et al., 2025). We present an alternative PEGDA MN hydrogel that incorporates GOx and the pH indicator bromocresol purple (BCP). In this design, GOx catalyzes the oxidation of glucose to gluconic acid, thus modulating the pH of the hydrogel. As glucose increases and pH decreases, BCP causes the patch to change color from purple to yellow, providing an intuitive read-out of blood glucose levels. Moreover, the biocompatible characteristics of PEGDA will allow for extended use with minimal health effects. Together, these findings highlight the potential of this MN hydrogel platform to transform glucose monitoring into a simple, cost-effective, accessible, and patient-friendly experience.

## MATERIALS AND METHODS

### Materials

Poly(ethylene glycol) diacrylate (PEGDA) (M_n_= 250 and M_n_= 700), lithium phenyl-2,4,6-trimethylbenzoylphosphinate (LAP), glucose oxidase (GOx, 100 U/mg), bromocresol purple sodium salt (BCP), and polydimethylsiloxane (PDMS) were all obtained from Millipore Sigma. A 365nm UV light source was purchased from Amazon (Everbeam 365nm 50W UV LED Black Light). Sheep polyclonal anti-GOx was obtained from Rockland Antibodies Online, and rabbit anti-sheep horseradish peroxidase conjugated was obtained from Abcam. All other reagents used were of analytical grade.

### Fabrication of PEGDA hydrogels

PEGDA hydrogels were created from varying concentrations of PEGDA 250 dissolved in ethanol (EtOH) or PEGDA 700 dissolved in PBS (pH ∼7). LAP photoinitiator was resuspended to 110mM in PBS and frozen in single-use aliquots at -80ºC. GOx was resuspended to 6000U/mL in PBS and flash-frozen using liquid nitrogen in single-use aliquots for long-term storage at -80ºC.

To make hydrogels, LAP was thawed and added to the precursor at a final concentration of 2.2mM. GOx was added to a final concentration of 5% v/v or 300 U/mL for relevant experiments **(Supp Fig. 1)**.

### Fabrication of hydrogel discs

Proof-of-concept experiments were performed using small hydrogels shaped into discs and incubated in PBS to allow for higher-throughput optimizations. Discs were formed by sandwiching 30µL of precursor between 2 RainX-treated glass microscope slides with a 1mm spacer **(Supp Fig. 2)** and crosslinking under UV light (365nm, 10-11.5mW/cm^2^) for 90 seconds.

### BCP acts as an effective pH indicator

To confirm that BCP would perform as an effective indicator for pH changes resulting from the reaction between GOx and glucose, we incubated GOx with 8mM glucose and 50µM BCP for 4 hours. The relationship between pH and BCP absorbance was established through a standard curve **(Supp Fig. 3)** and used to estimate the pH of the reaction in vitro, which decreased from 7 to 5 within 250 minutes over a reverse time course experiment **(Fig. 1A)**. This pH range allowed for a visible color change from purple to yellow within physiologically relevant glucose concentrations.

**Figure 1.**
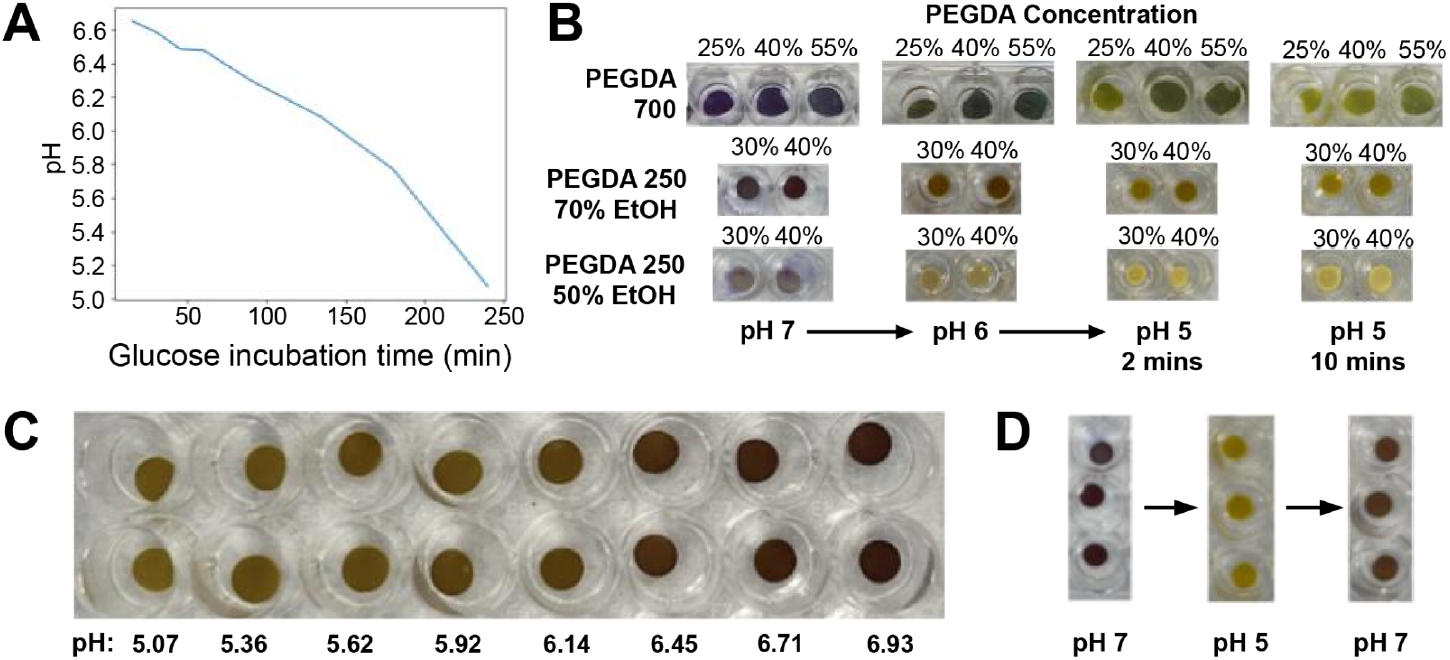
40% PEGDA 250 hydrogel discs demonstrate the most distinct and rapid color change. **(A)** pH of the GOx reaction decreases with time. **(B)** Hydrogels of varying composition with BCP are transferred from PBS at pH 7 to pH 6 and then to pH 5. Hydrogels were incubated in each pH for 10 minutes. **(C)** 40% PEGDA 250 hydrogels with 70% EtOH solvent are incubated in a range of pH from 5 to 7. **(D)** 40% PEGDA 250 hydrogels with 70% EtOH solvent are transferred from pH 7 to pH 5 and then back to pH 7.

### PEGDA composition testing

Next, we optimized hydrogel composition for BCP color response by testing PEGDA of molecular weights M_n_=250 and M_n_=700 at varying concentrations. To incorporate BCP into the hydrogels, PEGDA 250 or PEGDA 700 discs were swollen overnight in PBS with BCP at 500µM or 250µM respectively. PEGDA 700 was dissolved in PBS, whereas PEGDA 250 was dissolved in 50% or 70% EtOH.

In order to determine the hydrogel composition best suited for pH response, we incubated all gels in PBS at pH 7 before transferring them to pH 6 and then pH 5. PEGDA 700 gels did not show a clear color change until 10 minutes of incubation, while the PEGDA 250 / 70% EtOH condition demonstrated a color change within ∼2 minutes **(Fig. 1B)**. Of all tested conditions, 40% PEGDA 250 / 70% EtOH exhibited the fastest color change **(Supp Fig. 4A)** and a distinct color gradient across physiologically relevant pHs **(Fig. 1C)**. These hydrogels were also able to dynamically respond to repeated changes between incubation in pH 7 and pH 5 PBS via a clear color change from purple to yellow and back to purple **(Fig. 1D), (Supp Fig. 4B)**. Therefore, 40% PEGDA 250 in 70% EtOH was chosen to fabricate hydrogel discs.

### Characterization of hydrogel discs

We characterized the mechanical properties of the 40% PEGDA 250 hydrogel discs using oscillatory rheology. Amplitude sweeps confirmed a well-defined linear viscoelastic region (LVR) extending from 0.1% to approximately 1% shear strain. Within this range, the storage modulus (G′) exceeded the loss modulus (G″), indicating predominantly elastic, solid-like behavior for the initial compression of the patch to the skin. At 0.1% strain, the mean G′ was approximately 232±69 kPa while the mean G″ was about 132±43 kPa (n=5) **(Fig. 2A)**. Beyond ∼1% strain, both moduli declined monotonically, with a G′/G″ crossover at intermediate strain indicating network yielding and the onset of viscous flow. Frequency sweeps at 1% strain within the LVR showed that G″ exceeded G′ across the full angular frequency range of 0.1–100 rad/s, indicating viscous-dominant behavior under loading **(Fig. 2B-C)**. These data suggest that PEGDA 250 hydrogel discs possess reasonable mechanical strength and fidelity.

**Figure 2.**
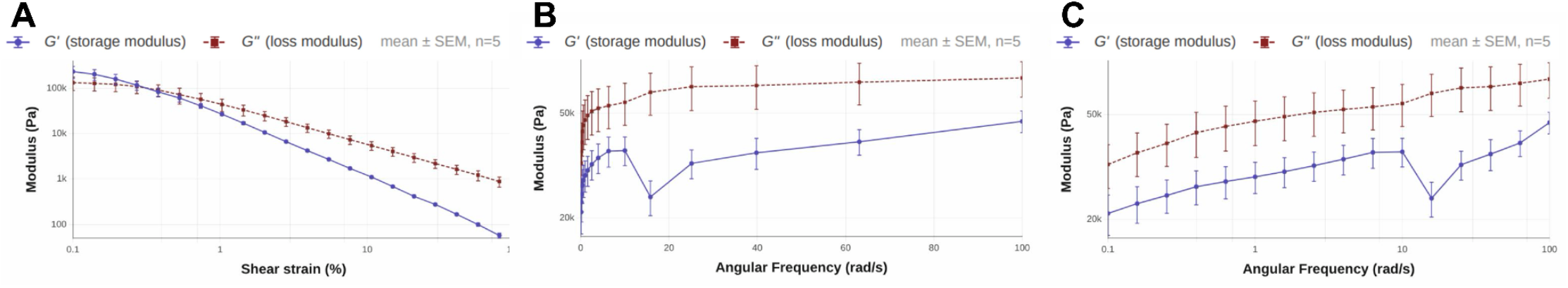
Mechanical properties of 40% PEGDA 250 measured by oscillatory shear rheometry. **(A)** Amplitude sweep: storage modulus (G′, filled circles, solid line) and loss modulus (G″, filled squares, dashed line) over shear strain (0.1 – 100%) at 1 rad/s. **(B)** Frequency sweep at 1% shear strain over 0.1 – 100 rad/s. **(C)** The frequency sweep data were linearized. Data reflect mean ± SEM with n = 5.

### Fabrication of hydrogel microneedles

After evaluating the mechanical properties of our gels, we began fabricating MN patches. We cast positive resin models of our MN patch made with light-based printing in PDMS to create negative molds. MNs were formed by adding 350µL of precursor to each negative PDMS MN mold, followed by vacuum desiccation for 20 minutes and then UV crosslinking **(Fig. 3A)**. We originally fabricated 8.1×8.1mm patches with an 11×11 array of 1mm long needles **(Supp Table 1)** but observed severe defects with scanning electron microscopy **(Supp Fig. 5)**. To consistently form needles, we increased the size of the patch to 16.2×16.2mm and the length of the needles to 2mm **(Supp Table 1)**.

**Figure 3.**
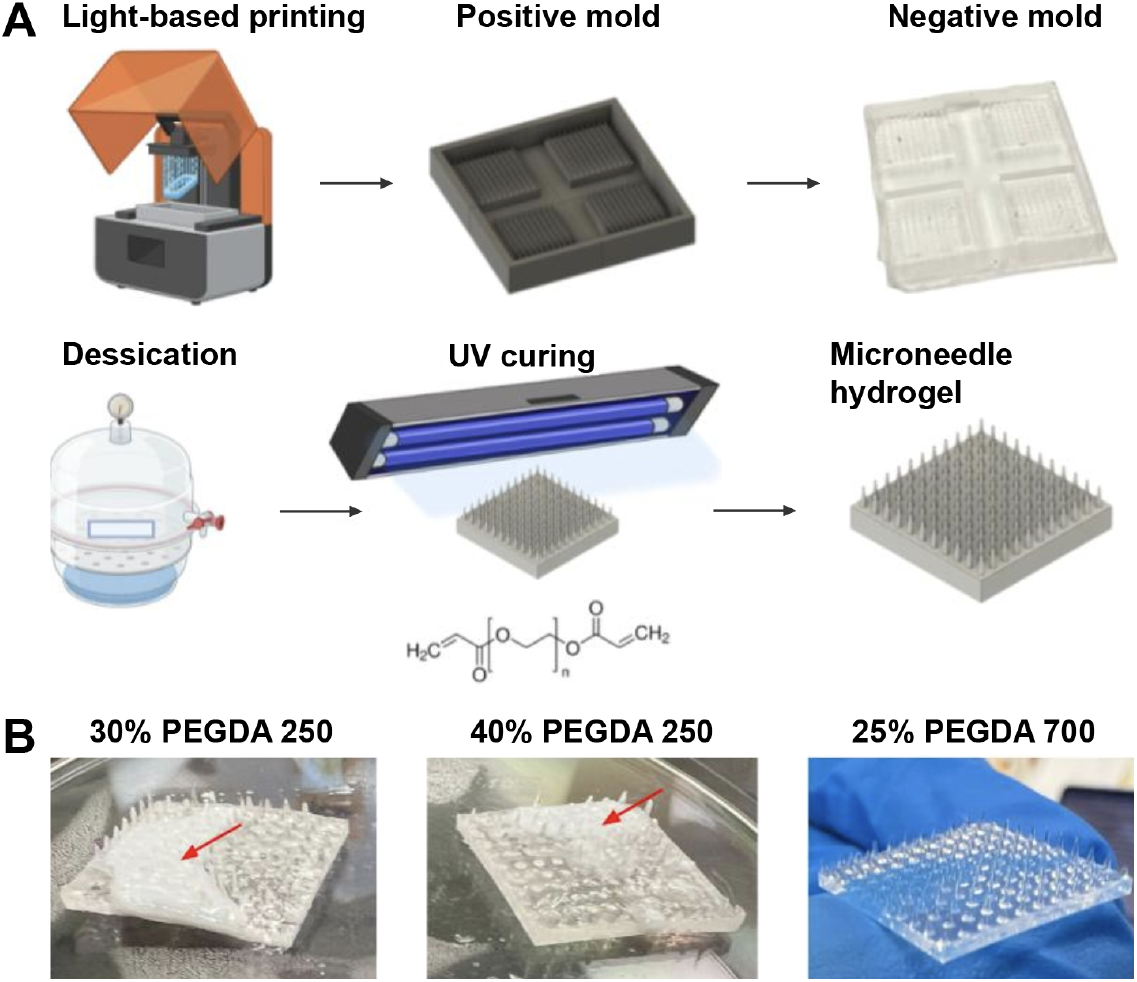
MN fabrication and composition testing. **(A)** Positive master molds were printed with ABS-like resin on a Halot-X1 3D printer, washed with isopropyl alcohol, dried for 5 days at 60 C, and used to make negative molds. MN hydrogels were prepared by 15 minutes of desiccation followed by 2 minutes of UV curing. **(B)** 30% and 40% PEGDA 250 hydrogels in EtOH and 25% PEGDA 700 hydrogels in PBS were desiccated into MN molds and crosslinked. PEGDA 250 gels formed patches of white and clear hydrogels marked by red arrows.

Unfortunately, we noticed that PEGDA 250 in EtOH would become insoluble following desiccation, leading to heterogeneous patches of white and clear hydrogel **(Fig. 3B)**. The low-pressure environment in the desiccator may have caused EtOH in the solvent to vaporize, reducing PEGDA 250 solubility in localized patches. In contrast, 25% PEGDA 700 hydrogels in PBS produced a uniform patch with clear, sharp MNs **(Fig. 3B)**. Therefore, we decided to fabricate MNs using 25% PEGDA 700.

## RESULTS

### Hydrogel discs loaded with GOx exhibit dynamic color change in response to glucose

Following confirmation of BCP as a viable indicator and the tolerance of GOx and BCP to UV radiation and EtOH **(Supp Fig. 6)**, we incorporated GOx into hydrogel discs to evaluate the efficacy of the GOx-glucose reaction within a hydrogel. Discs were incubated in PBS with glucose at 0, 54, 99, and 144 mg/dL. Control samples with no GOx remained purple, while samples with GOx exhibited an increase in yellow color with increasing glucose concentration **(Fig. 4A)**. To determine if the hydrogels could dynamically respond to a change in glucose concentration, we removed the discs from their initial incubation buffer after 280 minutes. Gels initially incubated in 0 and 144 mg/dL glucose and gels initially incubated in 54 and 99 mg/dL glucose were swapped and placed into fresh glucose solution of the other concentration. Gels were incubated for an additional 300 minutes and demonstrated clear reversal of colors to reach the expected equilibrium **(Fig. 4B)**.

**Figure 4.**
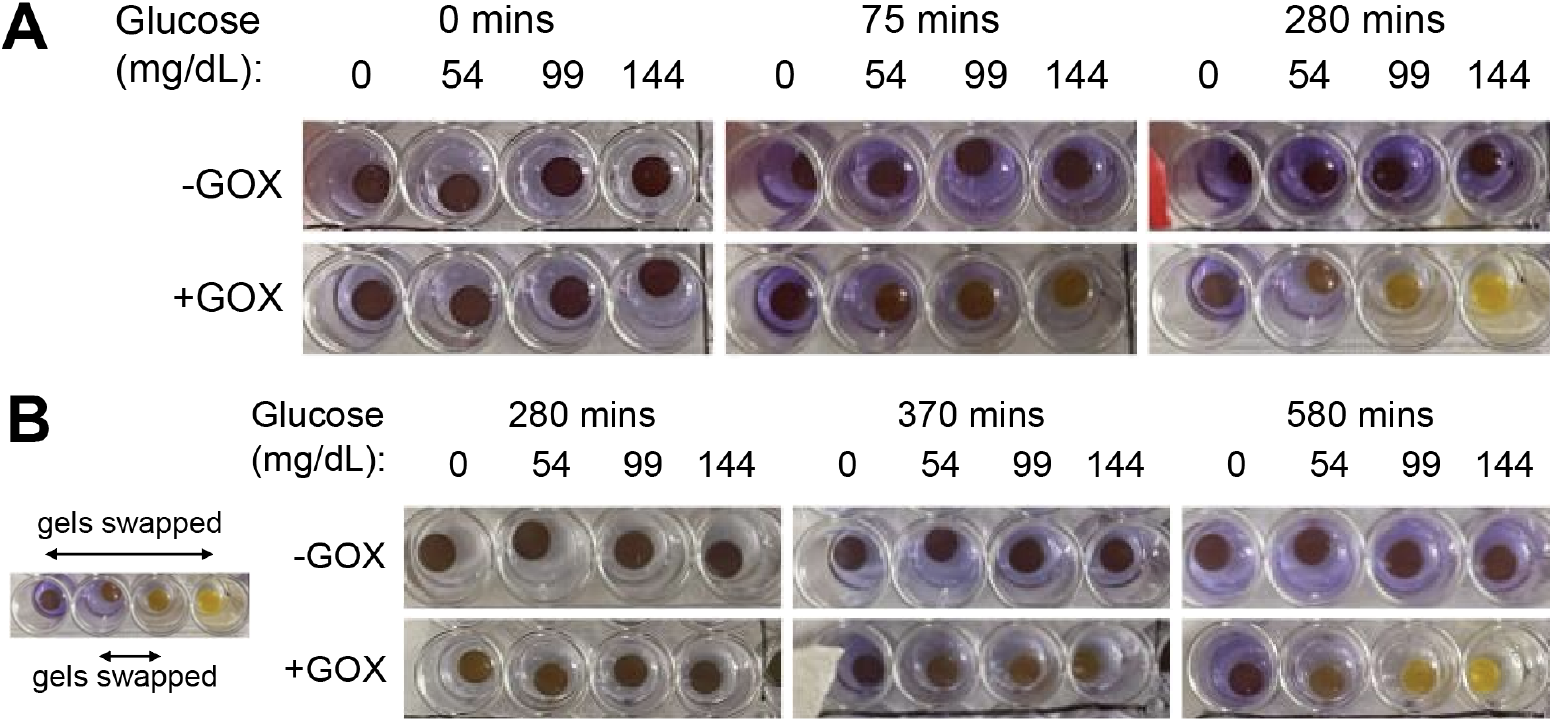
Glucose response of GOx-loaded hydrogels. **(A)** 40% PEGDA 250 gels loaded with GOx and BCP were incubated in glucose at 0, 54, 99, and 144mg/dL. **(B)** After 280 minutes, gels were swapped into fresh glucose solution of a different concentration and incubated for an additional 300 minutes.

### Quantitative analysis of glucose levels based on color change

Next, we developed a model relating the color of a hydrogel to glucose concentration using photographs captured by a mobile phone. Three hydrogel discs were incubated in each of 8 concentrations between 0 and 190 mg/dL glucose for 225 minutes before being photographed **(Fig. 5A)**. Mean pixel intensities for the red (R), green (G), and blue (B) channels were extracted from these photographs and independently fitted with third-degree polynomial regression models using all 24 data points **(Fig. 5B)**. Glucose concentration was predicted for 5 new hydrogel samples by identifying the point along the fitted 3D RGB curve that minimized Euclidean distance to the measured color coordinates of each unknown hydrogel. The model demonstrated strong predictive performance, with an RMSE of 8.6 mg/dL and R^2^ of 0.94 **(Fig. 5C)**. Predictions were consistent across the physiologically relevant glucose range of 63-171 mg/dL, with errors ranging from -9.0 to +14.6 mg/dL. The largest error occurred at the upper end of the standard range, which may reflect reduced color contrast between adjacent high-glucose concentrations as BCP approaches its yellow saturation point.

**Figure 5.**
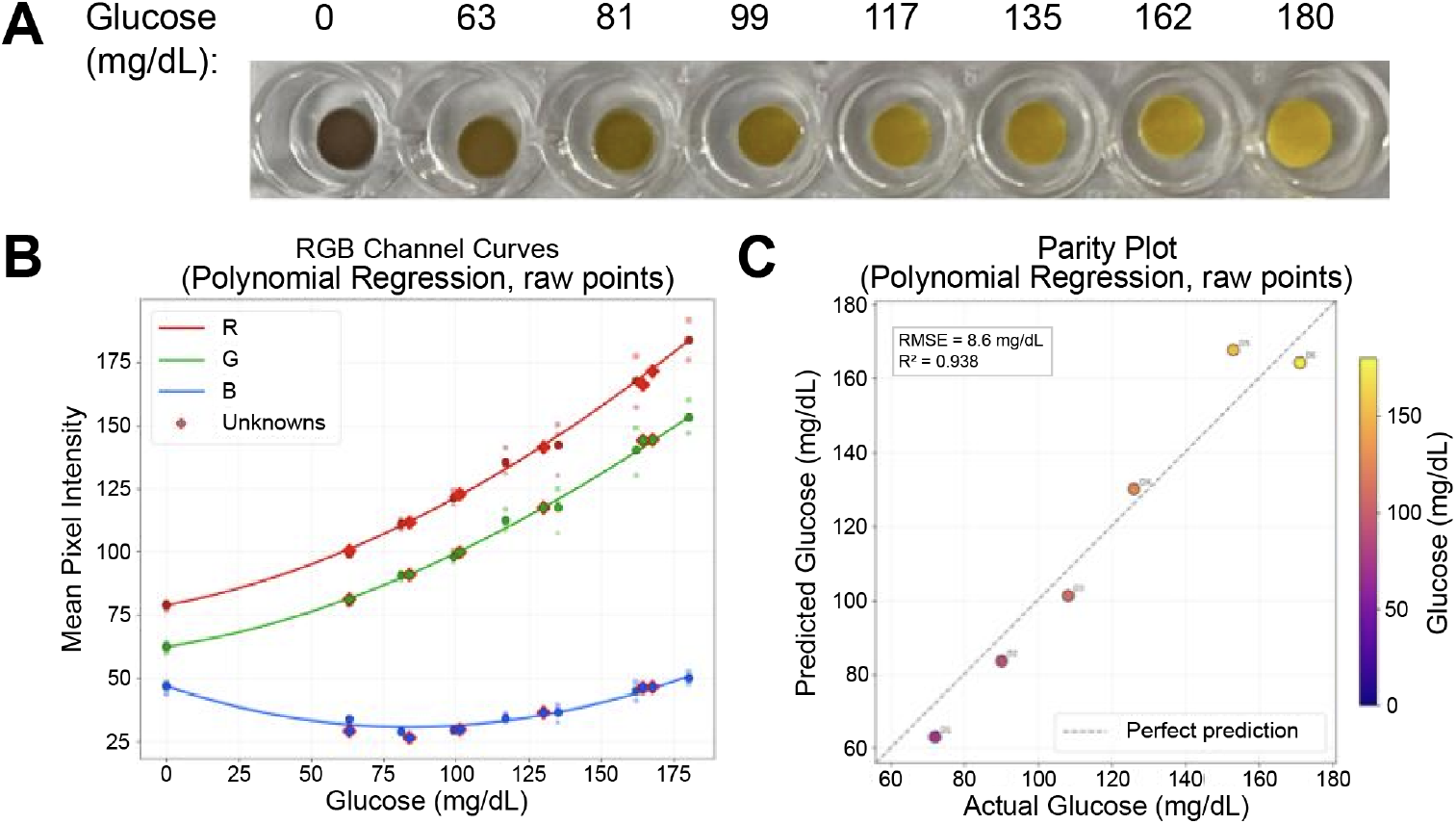
Glucose concentrations predicted from hydrogel color. (A) Images of hydrogels incubated in a range of glucose from 0-180mg/dL for 225 minutes (n=3). (B) Third-degree polynomial model predicting glucose concentrations from color. (C) Parity between actual and predicted glucose concentrations using images of 5 new hydrogels at a range of glucose concentrations (RMSE=8.59 mg/dL, R^2^ =0.938).

### Limited GOx diffusion observed out of hydrogel discs

Next, we evaluated the ability of these hydrogels to retain GOx through mesh size, which is essential for device function. Three replicates of each glucose buffer used previously to evaluate color change **(Fig. 4A)** were collected after 280 minutes of incubation with a GOx-loaded hydrogel. Replicates of each glucose concentration were pooled and stained for GOx on a Western blot **(Fig. 6A)**. Significant GOx was observed at all glucose concentrations. To quantify GOx diffusion, we collected the three replicates of the 0 mg/dL incubation buffer previously used to construct the glucose standard **(Fig. 5A)**. These samples were individually stained for GOx on a Western blot alongside a standard curve of serially diluted GOx. Comparing sample band intensity to the standards yielded an average diffusion of 6.5±2.32µg GOx per gel or 7.23±2.58% of the total 90µg of GOx loaded into each gel **(Fig. 6B-D)**. The hydrogel was able to retain ∼93% of the total GOx, allowing for continued catalysis of glucose oxidation.

**Figure 6.**
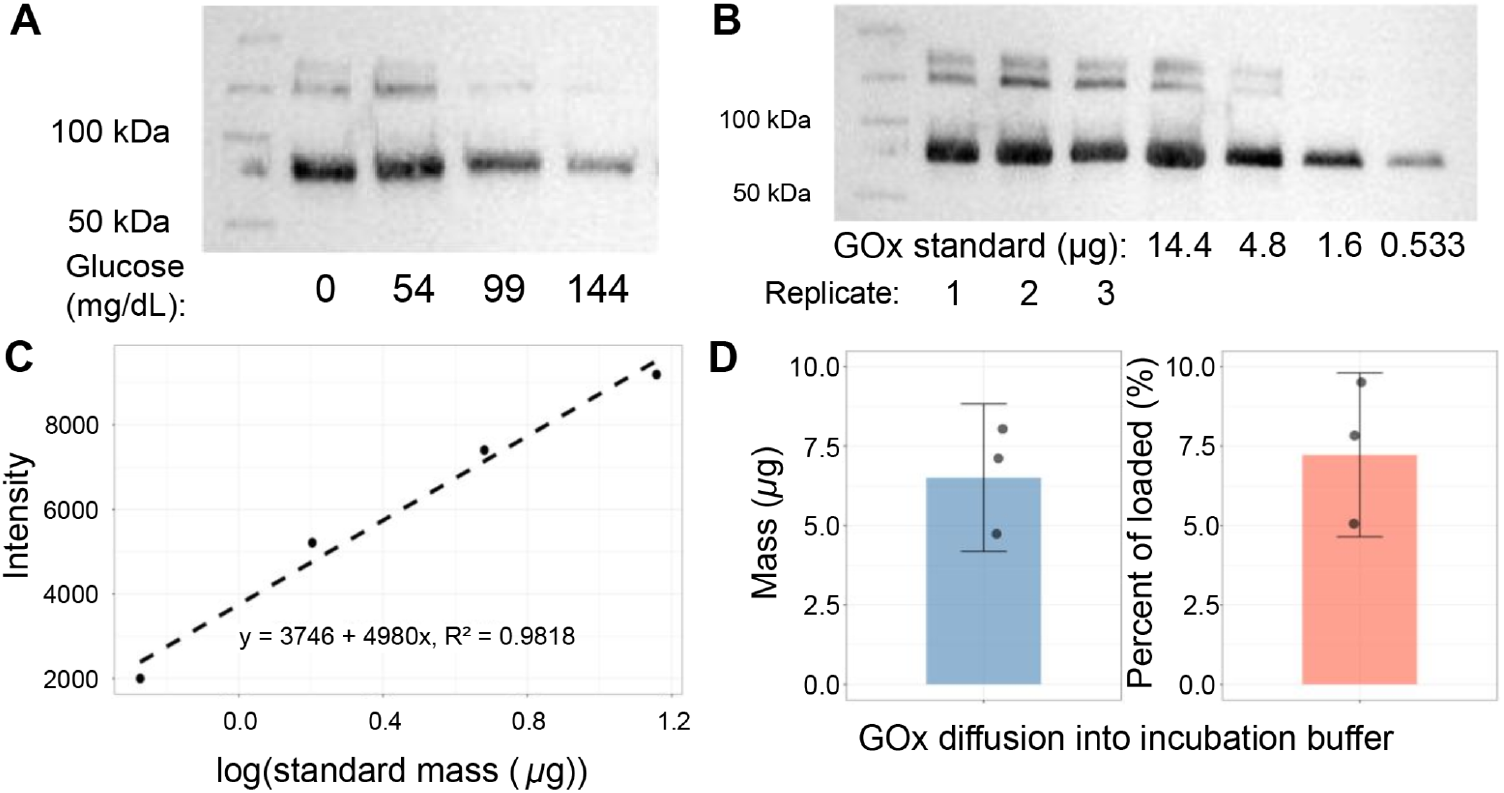
GOx diffusion out of hydrogel discs. **(A)** Glucose buffer incubated with GOx-loaded hydrogels stained for GOx at four glucose concentrations by Western blotting. **(B)** PBS incubated with GOx-loaded hydrogels (n=3) stained for GOx next to a GOx standard. (C) Band intensities of standards are used to construct a standard curve relating intensity to GOx quantity (R^2^ =0.9818). **(D)** GOx diffusion per gel is determined from the standard curve. Error bars represent standard deviation (n=3).

### MN hydrogel patches demonstrate distinct color change in glucose-loaded agarose

We next evaluated the ability of PEGDA 700 MN patches to change color in response to glucose. Patches loaded with GOx and swollen overnight in 250µM BCP were embedded into 1% agarose in PBS doped with glucose at 0, 75, and 150 mg/dL. Controls without MNs were included at each glucose concentration to test whether the MNs affected the rate of glucose diffusion. Controls were prepared by flipping the hydrogels upside down so that only the side without MNs was exposed to the agarose. After 24 hours of incubation, a clear color change from purple to yellow with increasing glucose concentration was observed **(Fig. 7)**.

**Figure 7.**
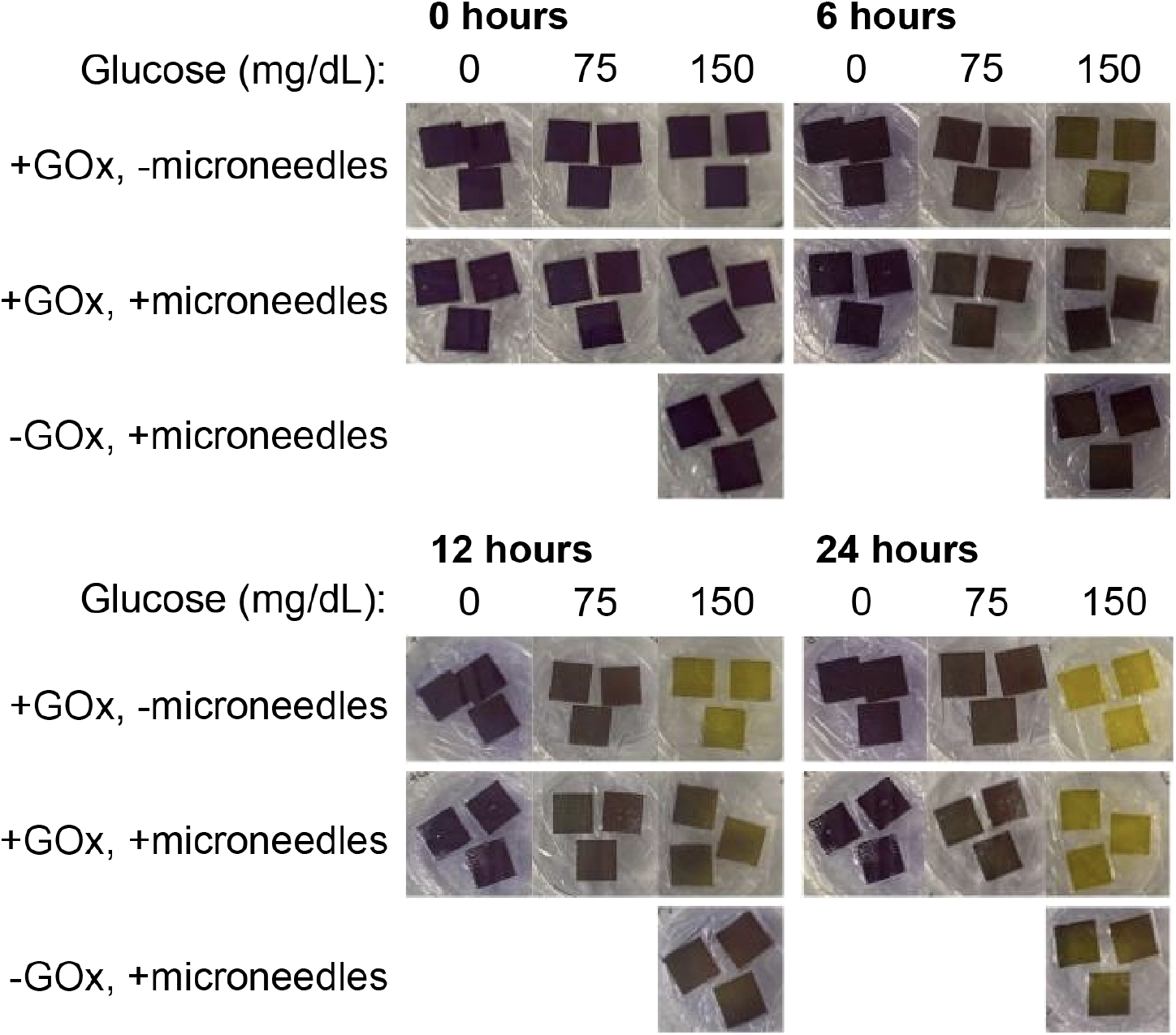
MN patches change color when embedded in agarose with glucose. Patches loaded with GOx and BCP are embedded in varying concentrations of glucose for up to 12 hours. No MN control patches were flipped so that only a flat interface contacted the agarose.

Surprisingly, hydrogels without MNs experienced a more rapid color change compared to hydrogels with MNs **(Fig. 7)**. We propose that the limited rigidity of the MNs led to imperfect embedding into the agarose. Needles that are bent or pressed out of shape may disrupt contact between the patch and agarose and reduce total contact area. Assuming that every MN was perfectly formed, with no blunting at the tips, 51.2% of the MN patch would need to be lifted away from the agarose to result in equal total surface area exposure compared to the non-MN control. Moreover, removing the MNs shortened the average distance required for the glucose to diffuse into the base of the patch, potentially decreasing the time until visible color change. Likely, a combination of improved contact with agarose and decreased diffusion distance resulted in a faster color change in the no MN conditions. Despite this unexpected result, the MNs would be essential to sample ISF in a transcutaneous application, where glucose is not present directly at the skin-device interface. We also observed a slight color change in patches created without GOx, suggesting that our gel-making process and re-use of the same molds could have led to enzyme contamination between samples.

### COMSOL diffusion and reaction kinetics modeling

To compare glucose transport and enzymatic reaction kinetics between hydrogels with and without MNs, a three-dimensional computational model was developed in COMSOL Multiphysics. The GOx-catalyzed oxidation of glucose to gluconolactone and hydrogen peroxide was modeled sequentially and followed by spontaneous hydrolysis of gluconolactone to gluconic acid. A simplified flat interface geometry was created to model the non-MN condition **(Fig. 8A)**. Glucose diffused progressively into the bulk hydrogel, establishing a concentration gradient with highest concentrations near the entry interface. Enzymatic reaction rates were greatest near this surface, producing gluconolactone that accumulated and diffused inward. Hydrolysis to gluconic acid reactions created a heterogeneous pH distribution with the greatest acid presence near the glucose entry point.

**Figure 8.**
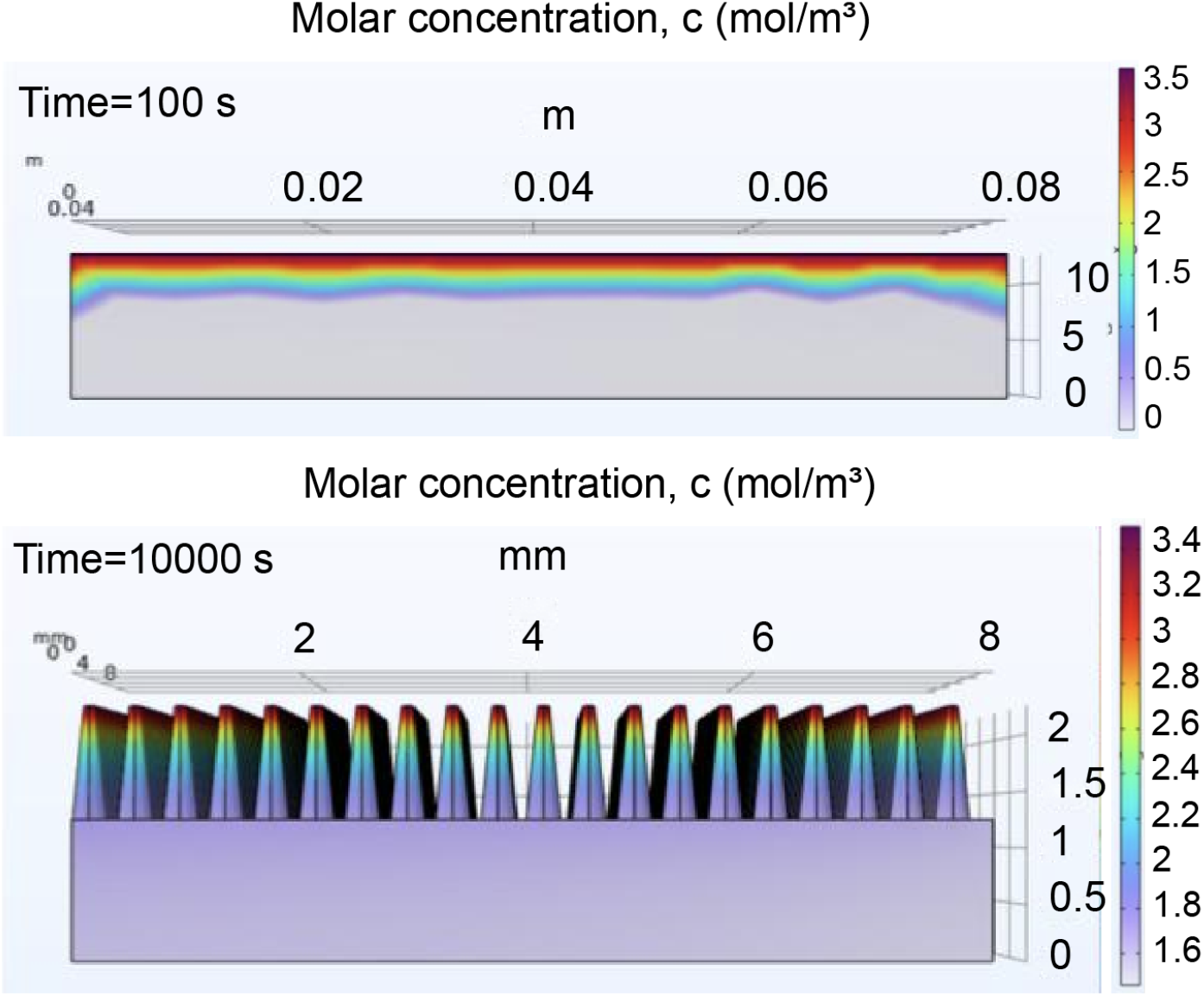
Glucose concentration gradients modeled in COMSOL. (A) Glucose gradient across the surface of the hydrogel at a time interval of 100 seconds indicates early signs of heterogeneity. (B) Full diffusion of glucose from the MN tips into the base of the patch is observed over a time interval of 10,000 seconds.

Subsequently, a digital twin of the MN hydrogel was created. Modeling diffusion of glucose from the MN tips into the base revealed that 2.7 hours were necessary for complete saturation of the hydrogel **(Fig. 8B)**, compared to 2.5 hours for the non-MN hydrogel. This corroborates the results of the previous experiment **(Fig. 7)**, with the non-MN hydrogel experiencing complete saturation at an earlier time point than the MN hydrogel. Even without considering imperfect agarose contact due to bent needles, the shorter average diffusion distance resulted in faster saturation of the non-MN model.

Although the non-MN hydrogel simulation resulted in faster complete glucose saturation than the MN hydrogel simulation, these models failed to account for the heterogeneous distribution of glucose in the skin. MN hydrogels must penetrate 250-1000 µm (Saifullah & Faraji Rad, 2023) into the skin in order to access and absorb ISF, thus validating the MN design **(Fig. 8A)**.

## DISCUSSION

Continuous glucose monitors have revolutionized diabetes care, providing patients with a minimally invasive and accurate method of tracking their blood glucose in real time. However, the use of expensive electronic components to detect hydrogen peroxide limits access to this technology to higher-income populations. We demonstrate a proof-of-concept, low-cost hydrogel device that detects gluconic acid instead of hydrogen peroxide, taking advantage of the pH change of the GOx reaction to report blood glucose concentration using BCP as a pH indicator.

First, we confirmed that pH changes from glucose oxidation fell within the relevant range for BCP colorimetric detection, validating BCP as an appropriate pH indicator for the GOx reaction. We then optimized PEGDA hydrogel composition to produce a distinct BCP-mediated color change across relevant pH levels **(Fig. 1)** and verified that the selected composition provided sufficient mechanical strength **(Fig. 2)**. Next, we incorporated GOx and BCP into hydrogel discs and observed a clear, dynamic colorimetric response to glucose **(Fig. 4)**. Notably, the hydrogel discs demonstrated reversible color changes in response to changing glucose concentrations, supporting the potential of this design to monitor fluctuating daily glucose levels. We demonstrated that photographs of these hydrogels could be used to construct a RGB standard that accurately predicts glucose concentration from gel color, achieving a RMSE of 8.6 mg/dL and R^2^ of 0.94 in a parity experiment **(Fig. 5)**. This principle could be applied to develop an application that allows users to quantitatively determine their glycemic state. Finally, we fabricated MN patches and demonstrated a colorimetric response when the patches were embedded in agarose gels to simulate transcutaneous application **(Fig. 7)**.

Although rheological and SEM data establish that PEGDA 250 arrays possess reasonable mechanical strength and fidelity, it is unclear if they are sufficiently stiff to puncture skin. Rheological testing of PEGDA 250 discs measured an elastic storage modulus of ∼232 kPa within the LVR **(Fig. 2)**, indicating that the hydrogel disc tested is substantially stiffer than the epidermis (Young’s modulus ∼0.1–1 MPa). Indeed, the hydrogel behaves as a viscoelastic fluid rather than a rigid elastic solid. This frequency-dependent behavior may be advantageous in reducing mechanical trauma at the needle–tissue contact. Needle stiffness under rapid loading is lower, meaning insertion speed and force should be tightly calibrated to remain within the elastic region for it to access ISF. However, we did not perform rheology testing on PEGDA 700 discs or the needles themselves, which likely differ significantly in their mechanical properties compared to PEGDA 250. PEGDA 700 MN hydrogels did not have sufficient strength to penetrate agarose without being damaged **(Fig. 7)** and are unlikely to puncture animal skin. Further work would continue to optimize the hydrogel composition, potentially exploring other polymers such as gelatin methylacryloyl and polyvinyl alcohol that are commonly used in MN fabrication. Changing the MN geometry could also improve their mechanical properties, as shorter and wider needles have been associated with increased mechanical strength (Zhang et al., 2025).

Another limitation of our design is the diffusion of BCP and enzyme out of the hydrogels. BCP was selected due to its sensitivity across the low pH range of the GOx reaction. However, this indicator is recognized as a mild skin irritant, with reported effects including itching and redness (PubChem, n.d.). Notable diffusion of BCP occurred in most experiments as seen by colored PBS incubation buffers and agarose **(Fig. 4; Fig. 7)**. Western blotting **(Fig. 6)** also showed significant diffusion of the enzyme, which may lead to immune reactions or other health issues. Future work will focus on functionalizing the PEGDA polymer backbone with BCP and GOx to ensure that both are retained within the hydrogel. Additionally, we plan to test our product using COMSOL simulations and empirical diffusion studies to assess potential exposure at the skin interface. If BCP contact with skin continues to present a health risk, we will work to identify alternative pH indicators that have established non-toxic effects on the body.

Our hydrogels also suffer from a slow response time to glucose. Although initial color responses can be observed within minutes **(Fig. 1)**, PEGDA 250 hydrogel discs require 280 minutes to reach equilibrium color changes, whereas PEGDA 700 microneedles were incubated for up to 24 hours **(Fig. 4; Fig. 7)**. The delayed response is governed by the combination of diffusion-limited transport and enzyme reaction kinetics. COMSOL modeling supports this interpretation, demonstrating that reactions are localized to the surface and estimating a 2.7 hour incubation period to allow for diffusion of glucose throughout the entire hydrogel for readable pH levels for BCP **(Fig. 8)**. Together, the enzymatic and transport limitations contribute to the observed lag in color change. To improve upon this, future work would involve optimizing the hydrogel structure to reduce crosslinking density, stabilizing enzyme loading through immobilization techniques, increasing the concentration of GOx in our hydrogels, or enhancing MN-tissue contact.

Despite these limitations, our results support a novel approach to monitor blood glucose through BCP-facilitated colorimetric indication of the GOx reaction. Further work to optimize hydrogel chemistry and functionalize the polymer network with BCP and GOx could result in a product suitable for animal testing and downstream clinical trials. Moreover, the continued development of a MN hydrogel digital twin using COMSOL would allow for further validation of experimental results. The fitted 3D RGB curve could also be further developed into a digital application for more accurate readings of colorimetric outputs. Altogether, the proposed electronic-free, highly cost-effective MN hydrogel design has the potential to expand access to CGM technology and, in turn, reduce the burden of diabetes-related complications in underserved populations.

## ACKNOWLEDGEMENTS

We thank Dr. Cameron Kim and Dr. Emma Chory, course instructors for BME 406/706: Biotechnology Design I/II, for their guidance throughout the project, and Aradhana, graduate teaching assistant for the course, for her mentorship on experimental design, troubleshooting, and laboratory technique. We also thank Margaret Gatongi, Senior Teaching Laboratory Administrator, and Melanie Tran, Research and Development Engineer, in the Duke University Department of Biomedical Engineering for their training and hands-on support in the teaching laboratory. We would also like to offer special thanks to Segura Lab members Connor Miksch for technical mentorship and Fernando Alamilla Munoz for assistance with the Halot-X1 3D printer.

## AUTHOR CONTRIBUTIONS

**AD:** Conceptualization, Investigation, Methodology, Validation, Visualization, Writing; contributed to project concept; involved in direct research activities, specifically focusing on enzyme and indicator validation, hydrogel composition optimization, and MN fabrication.

**HN:** Conceptualization, Investigation, Methodology, Writing; contributed to project concept; involved in direct research activities, specifically focusing on enzyme and indicator validation, hydrogel composition optimization, and MN fabrication.

**JD:** Conceptualization, Investigation, Project Administration, Visualization, Writing; contributed to project concept; involved in direct research activities, specifically reaction mechanism validation and MN fabrication; managed correspondence between team members and mentors.

**MD:** Conceptualization, Methodology, Formal Analysis, Software, Data Curation, Visualization, Writing; contributed to project concept; involved in formal analysis and processing, specifically focusing on colorimetric image quantification and RGB analysis.

**SS:** Conceptualization, COMSOL, Graphics, Model Analysis, Writing; contributed to project concept, involved in characterization of chemical kinetics for modeling and defining project mathematically for computational models.

**YL:** Conceptualization, Investigation, Methodology, Writing; contributed to the project concept; involved in direct research activities, specifically focusing on initial screening of hydrogel chemistries, material characterization via oscillatory rheology and SEM, and MN fabrication.

**EC:** Project administration, writing - review and editing; co-instructor for BME405L/406L: Biotechnology Design I/II.

**CK:** Funding acquisition, project administration, writing - review and editing; co-instructor for BME405L/406L: Biotechnology Design I/II.

## ETHICS STATEMENT

No institutional review board approval was necessary for this project

