## Supplementary Methods and Figures for "Hydrogel microneedle patch for low-cost, colorimetric, continuous glucose monitoring"

#### MN mold fabrication

A positive master mold was printed using ABS-like (Amazon, ELEGOO ABS-like 3D Printer Resin 3.0) on the Halot-X1 16k 3D printer (PioCreat, Cat# 1003040230). Following printing, the master mold was washed twice in isopropyl alcohol in a water bath sonicator, once for 3 minutes and again for 5 minutes. The master mold was then placed under UV light for 20 minutes and dried at 60°C for 5 days. Negative polydimethylsiloxane (PDMS) (Sylgard 184, Cat# 24236-10) molds were prepared using a 10:1 ratio of silicone elastomer to curing agent. The positive master molds were embedded in the PDMS in a 90 mm petri dish. The plate was placed in a dessicator for 15 minutes and then cured overnight at 60°C. After curing, a razor blade was used to cut out the negative molds and remove excess PDMS.

#### Hydrogel fabrication

PEGDA 250 (Millipore Sigma, Cat# 475629-100ML) and 700 (Millipore Sigma, Cat# 455008-100ML) hydrogels were created from precursors composed of varying concentrations of PEGDA 700 dissolved in 70% EtOH or PEGDA 250 dissolved in PBS. Lithium phenyl-2,4,6-trimethylbenzoylphosphinate (LAP) (Millipore Sigma, Cat# 900889-1G) photoinitiator was resuspended to 110mM in PBS and flash-frozen in single-use aliquots using liquid nitrogen for long-term storage at -80°C. For incorporation into hydrogels, LAP was thawed and added to a final concentration of 2.2mM. Discs were formed by sandwiching 30μL of precursor between 2 RainX-treated glass microscope slides with a 1mm spacer. Hydrogel discs were crosslinked under 365nm UV light (Amazon, Everbeam 365nm 50W UV LED Black Light) at an intensity of 10-11.5mW/$cm^2$ for 90 seconds. MNs were formed by adding 350μL of precursor to each negative PDMS MN mold, followed by vacuum desiccation within an HDMS chamber for 20 minutes before UV crosslinking. For the MN hydrogels embedded in agarose, the hydrogels were placed in a 90mm petri dish filled with 10mL of agarose and covered with Tegaderm transparent dressing film (Vitality Medical, Cat# 1622W) to prevent drying. To keep the hydrogels in place, a 35mm petri dish was placed inside the 90mm petri dish before adding a lid and wrapping in parafilm.

#### Enzyme and bromocresol purple incorporation

GOx (Millipore Sigma, Cat# G2133-10KU) was resuspended to 6000U/mL in PBS and flash-frozen in single-use aliquots using liquid nitrogen for long-term storage at -80°C. To incorporate into hydrogels, enzyme was thawed and added to precursor to a final concentration of 5%v/v or 300U/mL before crosslinking. BCP sodium salt (Millipore Sigma, Cat# 860891-5G) was loaded into crosslinked hydrogels through swelling. PEGDA 250 and PEGDA 700 hydrogels were swollen overnight by submersion in 500μM and 250μM BCP, respectively.

#### Western blotting and quantification

Samples containing GOx were denatured for 5 minutes at 95°C in Laemmli sample buffer (Bio-rad, 1610747) and beta-mercaptoEtOH before loading onto Bio-rad Mini-PROTEAN precast SDS-PAGE gels (Bio-rad, 4561024) running at 180V for 35 minutes. Proteins were transferred onto nitrocellulose using an Invitrogen Power Blotter Station. Membrane was stained for GOx using the ThermoFisher iBind Flex system with primary sheep anti-GOx (Rockland antibodies online, ABIN99713) and secondary rabbit anti-sheet HRP-conjugated (abcam, ab6747) antibodies. Chemilumunescence SuperSignal™ West Dura Extended Duration Substrate (Thermo Fisher, 34075) was incubated on the gel for 5 minutes before visualization using a Thermo Fisher iBright imager. Image band intensity was calculated using the built-in gel analysis tools of Fiji/ImageJ.

#### Plate reader

BCP concentration in buffers was determined by reading absorbance values at 595nm using a Victor NIVO Multimode Plate Reader and comparing to 2X serial dilution standards starting from 500μM BCP.

#### Rheology

The viscoelastic properties of UV-crosslinked PEGDA 700 hydrogels were characterized using an Anton Paar MCR 302 shear oscillatory rheometer equipped with an 8mm parallel plate geometry at 25°C. All measurements were performed in TruStrain™ correction mode. Amplitude sweeps were conducted at a fixed angular frequency of 1 rad/s over a shear strain range of 0.1–100% (n = 5) to identify the LVR and yield behavior. Frequency sweeps were subsequently performed at a fixed shear strain of 1% over an angular frequency range of 0.1–100 rad/s (n = 5). Storage modulus (G′) and loss modulus (G″) were recorded as functions of strain and frequency, respectively. Results are reported as mean ± SEM.

#### Scanning Electron Microscopy

Cross-sections of individual MNs were prepared using a Leica EM TIC 3x triple ion beam milling system (Leica Microsystems, Germany) with a high-energy argon-ion beam to produce smooth cross-sectional faces. Samples were subsequently sputter-coated with gold using the same ion beam milling system prior to imaging. SEM micrographs were acquired on a

Hitachi TM3030Plus tabletop SEM (Hitachi High-Tech, Japan) at an accelerating voltage of 15 kV in booster mode. Images were collected at low magnification (scale bar of 2mm) for array-level geometry and at higher magnification (scale bar of 500µm) for individual needle surface characterization. Needle length, area, and volume measurements were performed in ImageJ with scale bar calibration.

### Color-glucose quantification and prediction

Hydrogel color change was quantified from a single photograph of the entire well plate, captured with an iPhone 13 camera to ensure consistent lighting, distance, and imaging angle across all samples. Mean pixel intensity values for the red (R), green (G), and blue (B) channels were extracted from a fixed circular region of interest centered on each hydrogel disc using Fiji (ImageJ, NIH). Extracted mean RGB values were exported and analyzed using a custom Python script (Python 3.11; NumPy, SciPy).

To construct the glucose standard curve, all 24 raw replicate RGB measurements (3 replicates × 8 glucose concentrations spanning 0-180 mg/dL) were independently fitted with third-degree polynomial regression models for each channel using the numpy.polyfit function. The three channel-wise polynomial models collectively define a parametric curve through 3D RGB color space as a function of glucose concentration. Glucose concentration was predicted for unknown samples by identifying the point along this curve that minimized the Euclidean distance to the measured RGB coordinates of each unknown hydrogel, solved via bounded scalar minimization (scipy.optimize.minimize_scalar). Model accuracy was assessed by a parity plot comparing predicted versus actual glucose concentrations across six hydrogels at known glucose concentrations not used in standard curve construction, with performance reported as RMSE and $R^2$.

### COMSOL Modeling

A three-dimensional digital twin model was created in COMSOL. The geometry of the hydrogel was discretized using COMSOL's built in mesh generator, with a fine mesh refinement at the MN surfaces. The geometry of the patch had a 8.1x8.1 $mm^2$ base with a depth of 1.25mm. Each MN extending from the base had a conical shape with a diameter of 0.3mm on the base and 0.1mm on the tip with a depth of 1mm. Each MN was spaced 0.4mm from each other. Glucose was introduced into the hydrogel domain exclusively at the MN's tip with a concentration of 3.5mol/$m^3$. The initial concentration of glucose throughout the bulk of the hydrogel was set to zero, and oxygen was treated as a species with a constant concentration throughout with a physiological interstitial value of 0.2mol/$m^3$. All remaining species (gluconolactone, gluconic acid, and hydrogen peroxide) were initialized to zero concentration throughout the domain, and a no flux boundary condition was applied to all external surfaces. Drug transport throughout the 3D hydrogel domain was governed by the diffusion-reaction equation implemented through COMSOL's Transport of Diluted Species interface using Fick's 2nd Law. The velocity field was set to zero, and transport was assumed to be purely diffusion-driven. The effective diffusion coefficient of glucose within the hydrogel was described using a density-dependent expression.

$$\frac{\partial C_{H^+}}{\partial t} = D_{H^+}\nabla^2 C_{H^+} + R_{\mathrm{GOx}}$$

Equation 1: Fick's 2nd Law as the governing equation. $H^+$ represents protons.

The enzymatic oxidation of glucose by GOx was modeled as a two-way Michaelis-Menten reaction, and the spontaneous hydrolysis of gluconolactone to gluconic acid was modeled as a first-order reaction as user-defined variables. The coupled system of transient reaction-diffusion equations was solved using COMSOL's time-dependent solver through the Chemistry interface and coupled to the transport of Diluted Species interface via the Reactions node. A few challenges were encountered during COMSOL modeling. The Michaelis-Menten rate expression required entry as a user-defined reaction rate rather than a standard rate counter, and species variables required additional coupling to model parameters and equations. A circular variable dependency error occurred due to the additional couplings for the reaction interface, prompting a repeat COMSOL model set-up.

### Quantification and Statistical Analysis

All quantitative experiments were performed with a minimum of 2 technical replicates. Statistical analysis was only performed on experiments with a minimum of 3 replicates.

### Key Resources Table

| REAGENT or RESOURCE | SOURCE | IDENTIFIER |
|---|---|---|
| **Antibodies** | | |
| Sheep polyclonal anti-glucose oxidase | Rockland antibodies online | CAT#ABIN99713; RRID:AB_10781121 |
| Rabbit anti-sheep HRP conjugated | Abcam | CAT#ab6747; RRID:AB_955453 |
| **Chemicals, peptides, and recombinant proteins** | | |
| Glucose oxidase (GOx) from Aspergillus niger | Millipore Sigma | G2133 |
| Bromocresol purple sodium salt | Millipore Sigma | 860891-5G |
| Poly(ethylene glycol) diacrylate (Mn 250) | Millipore Sigma | 475629-100ML |
| Poly(ethylene glycol) diacrylate (Mn 700) | Millipore Sigma | 455008-100ML |
| Polydimethylsiloxane (PDMS) | Millipore Sigma | 761036-5EA |
| Lithium phenyl-2,4,6-trimethylbenzoylphosphinate | Millipore Sigma | 900889-1G |
| ELEGOO ABS-like 3D Printer Resin 3.0 | Amazon | https://www.amazon.com/ELEGOO-UV-Curing-Photopolymer-Toughness-Precision/dp/B0CTGZKS7X |
| **Critical commercial assays** | | |
| SuperSignal™ West Dura Extended Duration Substrate | Thermo Fisher | 34075 |
| iBind™ Flex Western Device | Thermo Fisher | SLF2000 |
| Power Blotter Station | Invitrogen | PB0010 |
| 7.5% Mini-PROTEAN® TGX™ Precast Protein Gels | Bio-rad | 4561024 |
| 4x Laemmli Sample Buffer | Bio-rad | 1610747 |
| **Other** | | |
| Everbeam 365nm 50W UV LED Black Light | Amazon | https://www.amazon.com/Everbeam-Black-Light-Flood-365nm/dp/B08635F9CX |

### SUPPLEMENTAL FIGURES

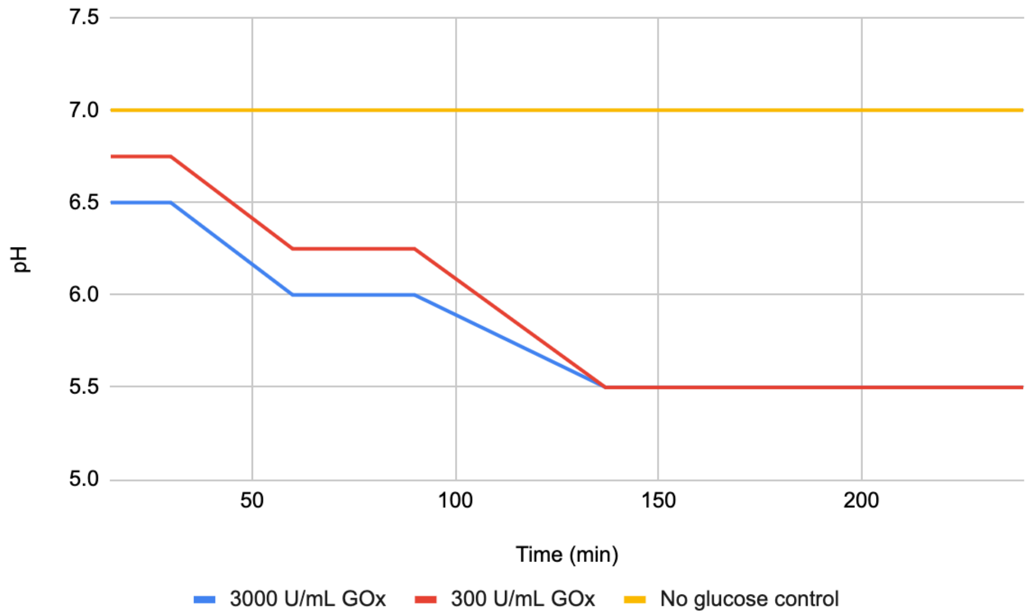


**Supplemental Figure 1) GOx quantity optimization.** 8mM glucose was incubated with 3000 or 300U/mL GOx for up to four hours. 1 of the initial 8 reactions for each GOx concentration was periodically sampled, and pH was estimated using a pH strip to find the pH of the GOx reaction over time. A control with 0mM glucose was included as well.

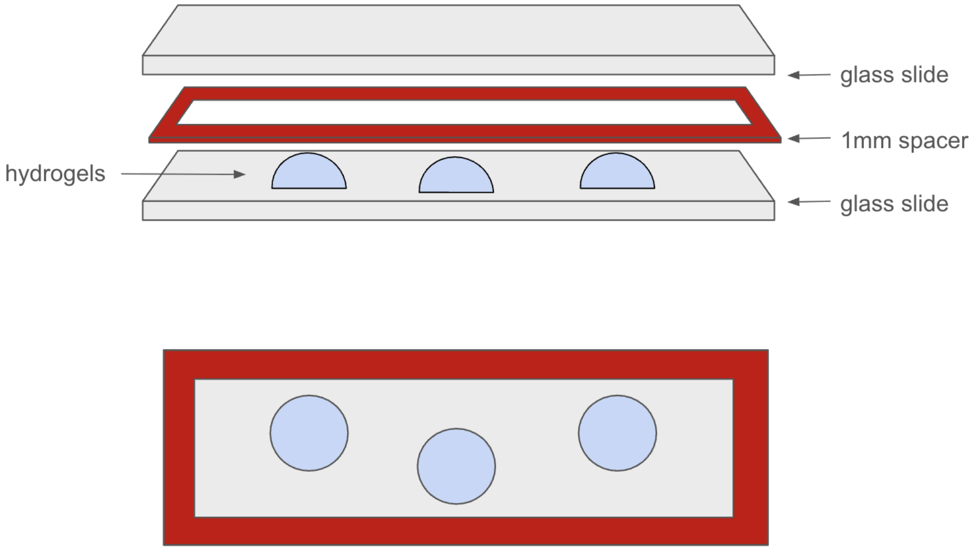


**Supplemental Figure 2) Hydrogel disc fabrication schematic.** For higher throughput experiments, hydrogel discs were fabricated by sandwiching 30 uL of precursor between two RainX-treated glass microscope slides with a 1mm spacer.

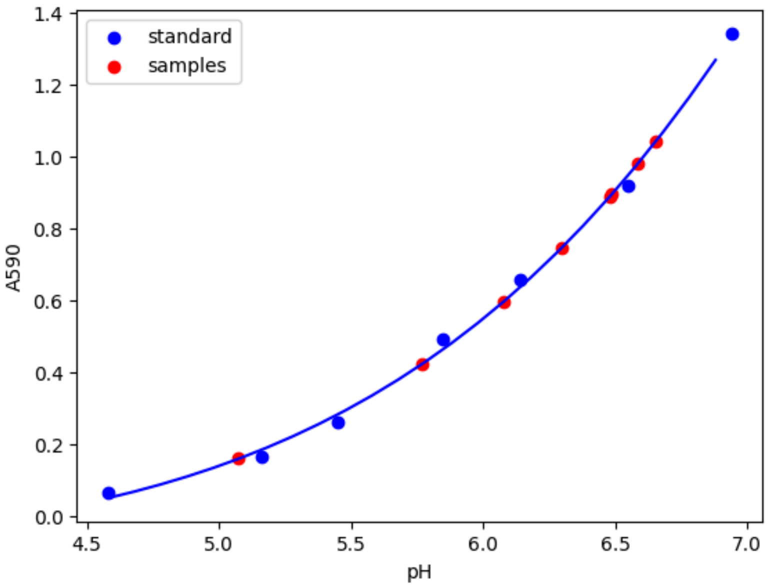


**Supplemental Figure 3) Standard curve between pH of 50μM BCP and A590.** PBS with 50μM BCP at varying pH values was measured for absorbance at 590nm (n=2). A 4-parameter logarithmic function is fit to the standards (blue) and used to predict the pH of the GOx reaction at different time points (red).

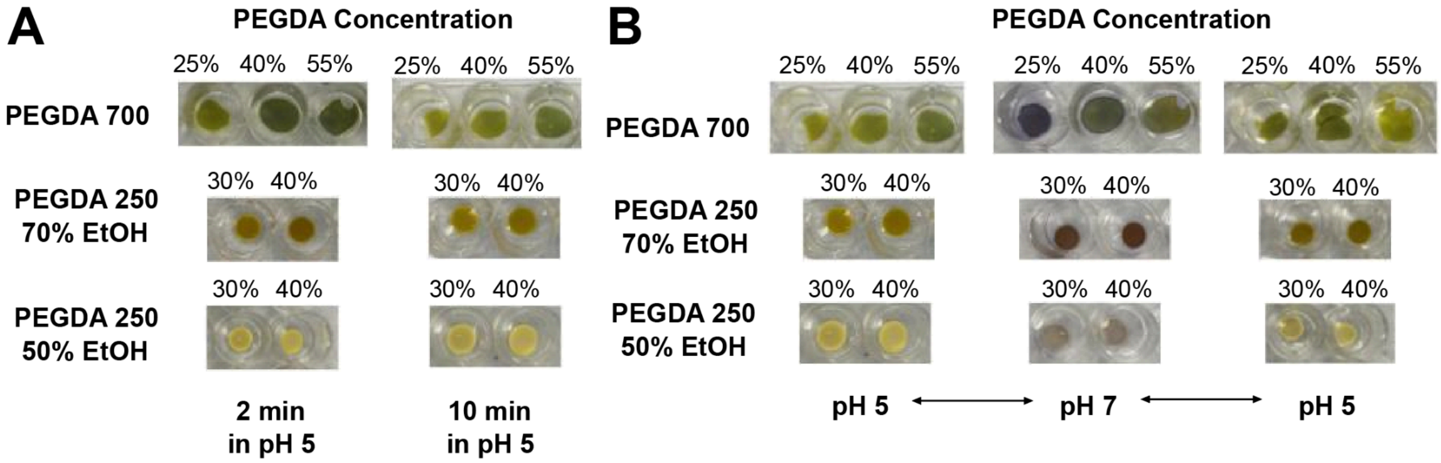


**Supplemental Figure 4) PEGDA composition testing. (A)** PEGDA 700 requires a longer incubation time for distinct color change than PEGDA 250. **(B)** All PEGDA compositions exhibited a dynamic color change response when transferred between PBS at pH 7 and pH 5.

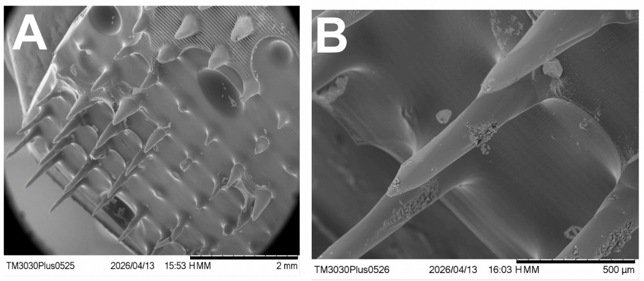


**Supplemental Figure 5) Electron microscopy of 8.1x8.1mm$^2$ PEGDA 250 MNs. (A)** Low-magnification micrograph of the MN patch showing defects in needle formation (scale bar of 2mm). **(B)** Higher-magnification image of individual MN shafts (scale bar of 500μm). Gold sputter coating applied prior to imaging. Images acquired using a Hitachi TM3030Plus SEM at 15kV.

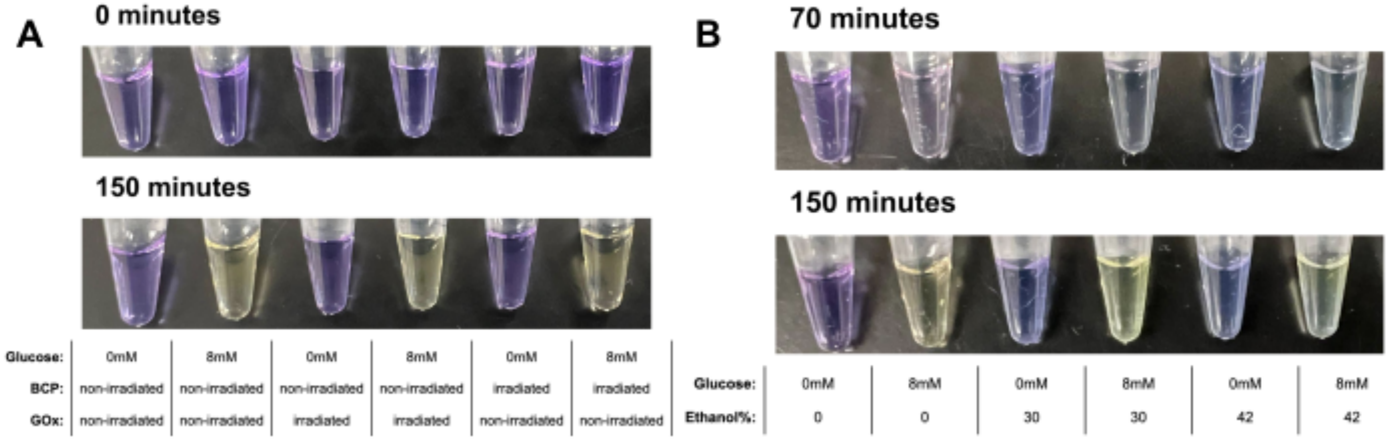


**Supplemental Figure 6) Radiation and EtOH tolerance of GOx and BCP. (A)** GOx and BCP both remain active after 90 seconds of ~10mW/cm$^2$ UV365nm radiation. **(B)** GOx and BCP remains active in EtOH concentrations of up to 42%

| | Base diameter (mm) | Tip diameter (mm) | MN height (mm) | Spacing between MNs (mm) | Array size | Total patch area (mm) | Thickness of base (mm) |
|---|---|---|---|---|---|---|---|
| **Original** | 0.3 | 0.1 | 1 | 0.4 | 11 x 11 array | 8.1 x 8.1 | 1.25 |
| **2x scaled** | 0.6 | 0.2 | 2 | 0.8 | 11 x 11 array | 16.2 x 16.2 | 1.25 |

**Supplemental Table 1) MN mold dimensions for both original and scaled up versions.**
